# The evaluation of miR-874 antagomiR and miR-146a toxicity in cardiomyocytes

**DOI:** 10.1101/2020.04.16.044925

**Authors:** Linlin Guo, Yue Zhang, Zhiwei Chen, Wenjing Dou, Jiamei Wu, Heli Xu, Hongyou Tan, Xiyan Wang

## Abstract

More and more studies have shown that microRNAs (miRNAs) play key roles in the treatment of heart failure. Studies have shown that miR-874 (miR-874) inhibitors can reduce H_2_O_2_-induced cardiomyocyte necrosis and may have a therapeutic effect on heart failure in terms of cardiomyocyte necrosis. Therefore, the purpose of this experiment is to study the other effects of miR-874 and miR-146a on the key pathways in cardiomyocytes and cardiac fibroblasts. In this study, the roles of miR-874 antagomiR and miR-146a in cardiomyocytes and cardiac fibroblasts were analyzed. we investigated the level of miR-874 expression in H9C2 cardiomyocytes after miR-874 antagomiR transfection and found that the miR-874 expression in the H9C2 cells transfected with miR-874 antagomiR group was significantly lower than that in the control group. The results showed that miR-874 was successfully knocked down by the transfection of miR-874 antagomiR. Our results demonstrated that miR-874 inhibition has no effect on the activity of Caspase-3/7 in cardiomyocytes. miR-874 antagomiR has no effect on the proliferation of cardiac fibroblasts, whereas it can inhibit the activity of caspase-8 in cardiac fibroblasts. In addition, the potential effect of miR-874 antagomiR on cardiac remodeling associated genes were examined. miR-874 antagomiR had no effect on SERCA2a mRNA level in H9C2 cells. In addition, miR-874 antagomiR were able to down-regulate the mRNA level of MMP9 and had no effect on MMP2 mRNA levels in H9C2 cells. Finally, The concentration of Ca^2+^ was measured using Fluo-4 NW Calcium Assay Kits following transfection of miRNAs or negative control in primary cardiomyocytes. miR-874 antagomiR was found to have no effect on Ca^2+^ concentration in cardiomyocytes. the concentration of Ca^2+^ in the cardiomyocytes transfected with miR-146a mimics was significantly lower than the mimic negative control group. In summary, miR-874 antagomiR and miR-146a may have a therapeutic effect on heart failure, but may also have side effects on heart failure treatment. Therefore, miR-874 antagomiR should be further studied to provide a basis for the development of drugs for heart failure.

## 1. Introduction

MicroRNA (miRNA) is a small non-coding RNA, composed of 19-25 nucleotides, involved in important biological processes, including growth, differentiation, apoptosis, and proliferation (Bartel DP et al., 2004;Yang D et al.,2017). A large number of studies have shown that miRNAs play key role in many heart diseases, such as myocardial infarction, cardiac hypertrophy, heart failure, myocardial fibrosis, arrhythmia, and hypertension (Yang B et al., 2007; Latronico MV et al., 2009), Recent studies have shown that miR-874 can inhibit the role of tumor cells in a variety of cancers (Jiang B et al., 2014; Nohata N et al., 2013; Nohata N et al., 2011; Kesanakurti D et al., 2013; Zhang X et al., 2014). There is little research on the role of miR-874 in heart failure. There is a study showing that miR-874 inhibitor can inhibit myocardial cell necrosis by targeting caspase-8, and miR-874 inhibitor provides new hope for the treatment of heart failure (Wang et al., 2013). There are many causes which lead to heart failure, including heart fibrosis and cardiomyocyte death. Cardiomyocyte death plays key role in heart failure,myocardial infarction and other diseases (Konstantinidis K et al., 2012). Cardiomyocyte apoptosis is one of the forms of cell death during the development of heart failure (Kajstura J et al., 1996). In addition, calcium overload causes cardiomyocytes to be damaged leading to heart failure (Arai M et al., 1994). Sarcoplasmic reticulum Ca^2+^-ATPase (SERCA2a) can regulate the uptake and release of Ca^2+^ in cardiomyocytes. Animal studies show that SERCA2a activity is significantly decreased in cardiomyocytes with heart failure and accompanied by Ca^2+^ transport disorders, sarcoplasmic reticulum intake of Ca^2+^ decreased and cytoplasmic Ca^2+^ aggregation during diastole, affecting the diastolic function of the heart, while the release of calcium decreased and cytoplasmic Ca^2+^ decreased during a systole, the corresponding decline in systolic function (Hu ST et al., 2010). Myocardial fibrosis is an important part of the process of heart failure. Increased cardiac fibrosis can cause decreased cardiac tissue contractility, eventually leading to heart failure (Dobaczewski M et al., 2012; Yarbrough WM et al., 2012; Kong P et al., 2014; Spinale FG et al., 2013). Cardiomyocyte proliferation and apoptosis are the direct cause of myocardial fibrosis. (Ma J et al., 2017) found that angiotensin II induced myocardial fibroblast proliferation promotes myocardial fibrosis. The matrix metalloproteinases (MMPs) are also important factors leading to myocardial fibrosis, especially MMP2 and MMP9. (Han CK et al., 2017). Induced myocardial fibrosis using lipopolysaccharide (LPS), and then gelatin zymography and Western blotting were used to detect MMP2 and MMP9 activity and protein level in H_9_C_2_ cells, It was found that the activity of MMP2 and the expression of MMP2 and MMP9 were increased in H_9_C_2_ cells after LPS treatment, the activity and content of MMP2 and MMP9 were the indicators of myocardial fibrosis. In summary, we investigated cardiomyocytes apoptosis, myocardial fibroblast proliferation and apoptosis, SERCA2a mRNA levels and Ca^2+^ concentration in cardiomyocytes, MMP2 and MMP9 mRNA levels in the study and used them as indicators to study the role of miR-874 inhibitors in the treatment of heart failure.

miR-146a has been considered as critical cardioprotective microRNAs strongly related to the pathogenesis of cardiac disorders,increased expression of miR-146a attenuated I/R induced myocardial apoptosis and promotes neonatal rat cardiomyocytes viability3.cardiosphere-derived cells (CDCs) are promising tools for cardiac cell therapy in heart failur patients,which have already been used in a phase I/II clinical trial to stimulate regeneration, angiogenesis, and functional improvement in the infarcted myocardium in both animals and humans. miR-146a was 262-fold higher in cardiosphere-derived cells (CDCs) exosomes than in normal human dermal fibroblasts (NHDF) exosomes; Furthermore,CDC exosomes might act through miR-146a transfer.

## 2. Materials and methods

### 2.1. H_9_C_2_ Cell Culture

Rat H_9_C_2_ cells were purchased from Shanghai Fumeng Inc and maintained in medium composed of Dulbecco’s modified Eagle’s medium (DMEM) high glucose (Wisent,Canada) supplemented with 10% fetal bovine serum (FBS),1% penicillin-streptomycin, and grown at 37°C in a humid atmosphere of 5% CO2/95% air until they reached 70-80% confluence.

The H_9_C_2_ cells frozen in liquid nitrogen were recovered and cultured in Dulbecco’s modified Eagle’s medium (DMEM) containing 10% fetal bovine serum. When the cell confluence was greater than 70%, the cells were passaged by routine method, and the cells in good growth condition were tested.

For in vitro overexpression studies, cells were transfected with miR-874 or a scramble miRNA mimics molecule for 48 h with RNAimax(Life Technologies) in OPTI-MEM reduced serum medium following the manufacturer’s recommendations.

### 2.2. Primary Cardiomyocytes And Cardiac Fibroblasts Culture

To obtain neonatal rat cardiomyocytes, 1- to 2-day-old Sprague-Dawley rats were decapitated and their hearts removed. Hearts were digested with a collagenase solution (Collagenase Type I, Life Technologies) followed by differential plating. Cells were plated at a density of 1×10^5^ cells/well in Ninety six-well plates, and cultured overnight in plating medium. Ara C was added to suppress the growth of the remaining fibroblasts.

### 2.3. Ethics statement

All the animal experiments and the animal welfare were in accordance with the Guide for the Care and Use of Laboratory Animal by International Committees. All procedures were approved by the ethics committee of Shen Yang Medical College, after treatment, RNAs were extracted from cardiac cells as described below.

### 2.4. Cell Transfection

The cardiac fibroblasts or H_9_C_2_ cells in good growth-condition were cell counted and adjusted to the appropriate concentration for seeding in the 96-well plates or 60mm dishes. Transfection experiments were carried out the next day when the cell confluence was about 60%. For primary cardiomyocytes, only the happy cardiomyocytes which can beat were chosen to do the transfection. The cells were transfected with 100 nM miR-874 inhibitor, negative control,miR-146a mimics or mimics control respectively according to the instruction of the transfection reagent RNAiMax. The cells were incubated in humidified air (5% CO_2_) at 37°C to prepare for subsequent experiments.

### 2.5. miRNA RT-PCR

Total RNA was extracted and purified using Trizol lysate and miRNeasy Mini Kit 48 h after transfection. The miRNA was reversely transcribed into cDNA using the Hairpin-it**™** miRNA and U6 snRNA normalization RT-PCR Quantition Kit, and U6 was used as the internal reference. miR-874 level was quantified after transfection of miR-874 antagomiR in H_9_C_2_ cells using ABI 7300 Real-time PCR system.

### 2.6. Cardiac Fibroblast Proliferation Assay

CellTiter-Fluor™ Cell Viability Assay was used to determine Cell proliferation 48h after transfection. The CellTiter-Fluor™ Cell Viability Assay reagents were mixed according to the manufacturer’s instructions. The excess medium was discarded and 50 μl of the final volume was retained in the 96-well plates. The 10 μl of CellTiter-Fluor ™ mixture was added to the medium of the cell and then the plate was shaken on a horizontal shaker for 30s and incubated for 30 min at 37 °C. Finally, the fluorescence was measured at 400_EX_ / 505_EM_ using the Biotek cyctation 5 cell imaging microplate reader. Fluorescence intensity reflects the number of viable cells, and then the effect of miR-874 antagomiR on the proliferation of cardiac fibroblasts was determined.

### 2.7. Cell Apoptosis Assay

The cells were starved overnight with medium containing 1% serum before transfection, and the CellTiter-Fluor™ Cell Viability Assay was used to detect cell number as internal controls 24 hours after transfection (see above). Apoptosis was measured by Caspase-Glo® Assay (including Caspase-Glo® 8 Assay detection of cardiac fibroblast apoptosis and Caspase-Glo® 3/7 Assay detection of cardiomyocyte apoptosis). The Caspase-Glo® Assay was mixed according to the instructions. 50 μl of the mixture was added to each well in 96-well plates. The plates were shaken for 30s on a horizontal shaker and incubated for 30 min at room temperature. Then Biotech cyctation 5 cell imaging microplate reader was used to detect the bioluminescence.The bioluminescence signal represented cysteine aspastic acid-specific protease (Caspase) activity in cell, reflecting the extent of myocardial fibroblast apoptosis.

### 2.8. qRT-PCR

Total RNA was extracted and purified using Trizol lysate and miRNeasy Mini Kit 48h after transfection of H_9_C_2_ cells. Total RNA was reverse transcribed into cDNA using High Capacity cDNA Reverse Transcription Kits. The mRNA levels of SERCA2a were detected by TaqMan® Gene Expression Master Mix and the Dmd was used as the internal reference. The mRNA levels of MMP9 and MMP2 were measured with Go Taq® qPCR Master Mix in the samples and the GAPDH was used as the internal reference. Specific primers are as follows:

Rat MMP2-F, 5’-CTTGCTGGTGGCCACATTC-3 ‘,

Rat MMP2-R, 5’-CTCATTCCCTGCGAAGAACAC-3 ‘;

Rat MMP9-F, 5’-CGCTCATGTACCCCATGTATCA-3 ‘,

Rat MMP9-R, 5’-TCAGGTTTAGAGCCACGACCAT-3 ‘;

Rat GAPDH-F, 5’-GGTGGACCTCATGGCCTACA-3 ‘

Rat GAPDH-R, 5’-CAGCAACTGAGGGCCTCTCT-3 ‘.

Real-time PCR was used to quantify mRNA expression and the amount of RNA was normalized to U6 expression. For miRNA analysis, total RNA was isolated using the mirVana miRNA isolation kit (Life Technologies). The levels of miRNAs were determined using miRNA specific Taqman probes from the Taqman MicroRNA Assay (Life Technologies). The relative amounts of miRNA were normalized to U6 expression. The qRT-PCR was performed with an ABI 7300 Real-time PCR system.

### 2.9. Detection of Ca^2+^ Concentration in Cardiomyocytes

After transfection for 48 h, Fluo-4 NW Calcium Assay Kits was loaded into the cardiomyocytes as calcium probe, Incubated at 37°C for 30 min and at room temperature for 30 min, Fluorescence intensity(F) was measured at 494_EX_ / 516_EM_ using Biotek cyctation 5 cell imaging microplate reader. After the Triton X-100(0.1%) was added and incubated at room temperature for 30 min, the maximum fluorescence intensity (Fmax) was measured. After the EGTA (5 mmol / L) was added and incubated at room temperature for 30 min, the minimum fluorescence intensity (Fmin) was measured. Finally, using the formula to calculate the concentration of calcium ions in cardiomyocytes: ([Ca^2+^] i = KD×(F-Fmin) / (Fmax-F), KD = 360nmol / L)

### 2.10. Statistical Analysis

Statistical analyses were carried out using independent t-test. P-values less than 0.05 were considered significant difference.

Results are expressed as the mean±s.d. of three independent experiments for *in vitro* studies, each consisting of three culture plates (n=9). Significant differences were established by either the Student’s t-test or one-way ANOVA, according to the number of groups compared.

## 3. Results

### 3.1. Cell Description

As shown, H_9_C_2_ cells were spindle-shaped and grew well (Fig.1a). After 3 days of culture, the cardiomyocytes had stretched out the pseudopodia, and the cells were connected into pieces and had regular spontaneous pulsation (Fig.1b). Cardiac fibroblasts were irregularly or elliptic, and became spindle-shaped, and the volume and nuclei were larger, no spontaneous pulsation occurred (Fig.1c).

**Fig. 1.**
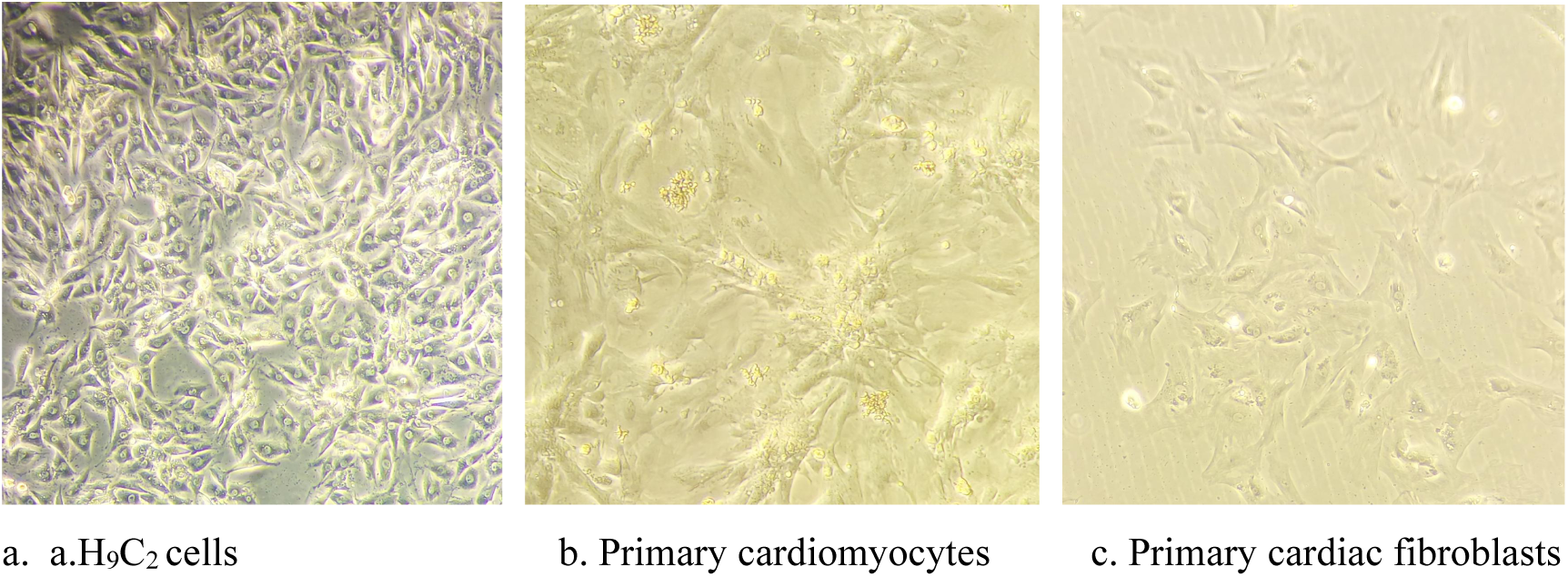
Cells are cultured in medium containing 10% serum.

### 3.2. Identification of downregulated miR-874 expression in cardiomyocytes following transfection of miR-874 antagomiR

In order to evaluate that miR-874 antagomiR was able to reduce the level of miR-874 expression in H_9_C_2_ cells, Quantitative RT–PCR was employed to detect the expression of miR-874 relative to U6 in cells. H_9_C_2_ cells were plated into 60mm culture dishes and transfected with 100nM miR-874 antagomiR or negative control. The cells were lysed and collected 48h after transfection. Total RNA was extracted with Trizol and reverse transcription was used to produce cDNA and miRNA RT-PCR was used to detect the level of miR-874 in H_9_C_2_ cells.

The results demonstrated that the miR-874 expression levels were significantly lower in miR-874 antagomiR transfected cells in comparison with scramble antagomiR control transfected cells (P<0.01,**Fig.2**), indicating that the miR-874 antagomiR successfully knocked down the level of miR-874 in H_9_C_2_ cells.

**Fig. 2.**
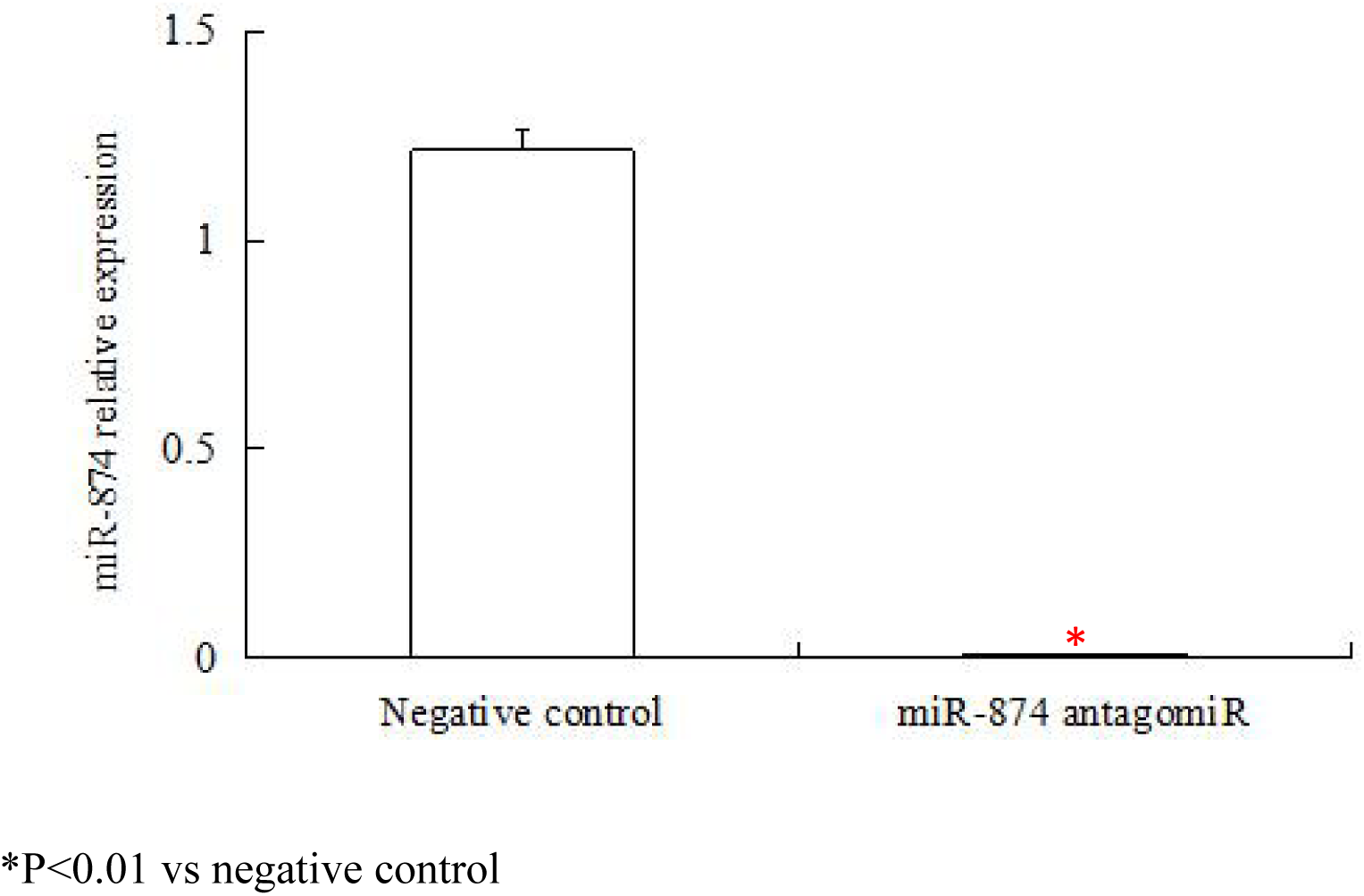
Transfection of H_9_C_2_ cells with miR-874-5p inhibitor markedly reduced miR-874-5p level.miR-874 antagomiR knocked down miR-874 expression levels in H_9_C_2_ cell. Quantitative Real-time RT–PCR showed that miR-874 expression in miR-874 transfected cells was significantly lower than that of negative control miRNA transfected cells. U6 snRNA was used as an internal control. *P<0.01 vs negative control

### 3.3. miR-874 antagomiR does not have effect on the caspase 3/7 activity

Cardiomyocyte apoptosis has been found clinically relevant to congestive and ischemic heart failure and may contribute to the development and progression of cardiac dysfunction and dilated cardiomyopathy. Caspase-3 has been demonstrated to be primary executioner caspase and key mediators of apoptosis in mammalian cells. However, it is not yet clear whether caspase-3 is a target of miR-874 and the molecular regulation of caspase-3 by miR-874 in H_9_C_2_ cells remains to be elucidated. (Because cardiomyocyte apoptosis is one of the causes of heart failure), in order to study the effect of miR-874 antagomiR on cardiomyocyte apoptosis, we performed Caspase-Glo® 3/7 Assay to detect the apoptosis of cardiac fibroblasts, the CellTiter-Fluor™ Cell Viability Assay was employed to detect the cell viability of cardiac fibroblasts. We added the two reagents in the same well to ensure that the activity of Caspase-3/7 is normalized to the corresponding number of cells, the number of cells corresponding to fluorescence activity and the activity of Caspase-3/7 corresponding to bioluminescence detection of the cell apoptosis, because the fluorescence and bioluminescence do not interfere with each other, so it can be superimposed. There is a increase in caspase3 activity in miR-874 knockdown cells, whereas there is no significant difference between the activity of Caspase-3/7 normalized to the number of cells in the miR-874 antagomiR group and the control group as shown in Fig.3.

**Fig. 3.**
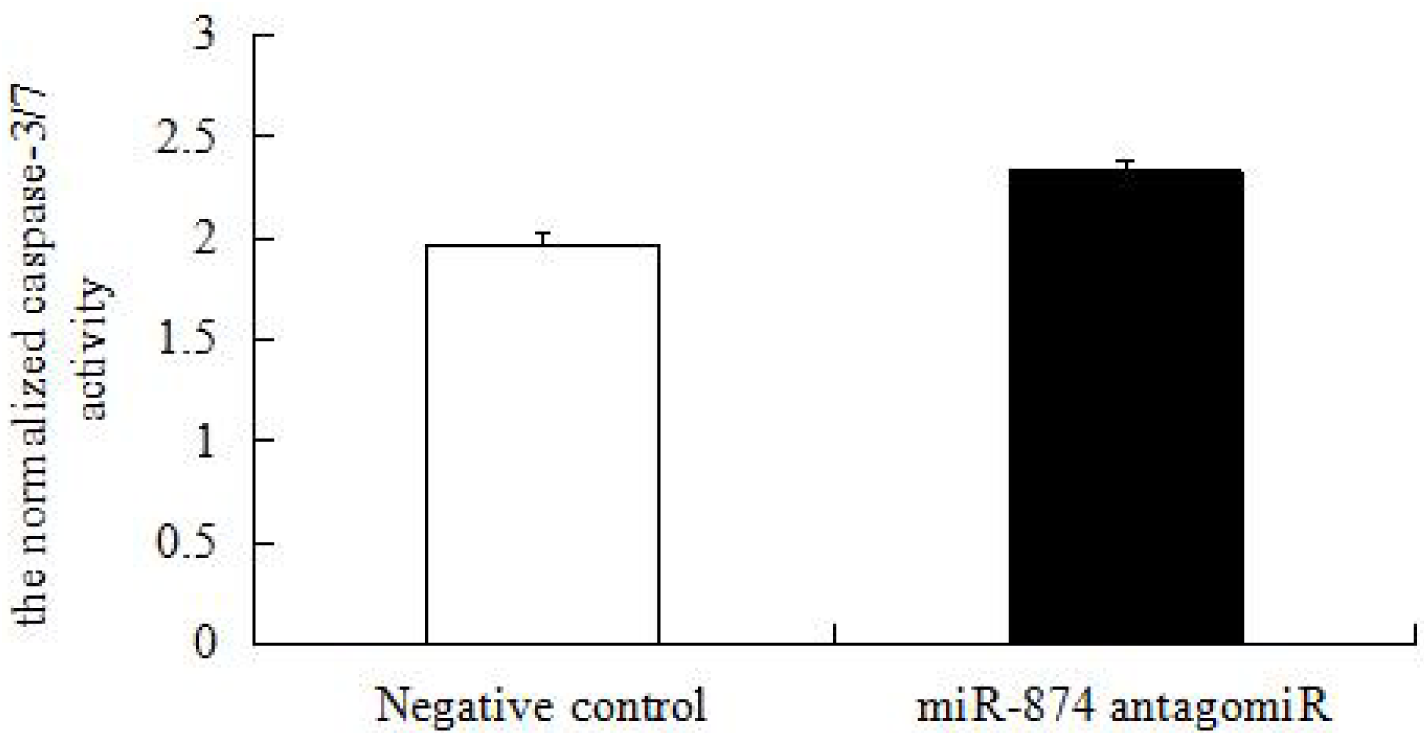
miR-874 antagomiR does not have effect on the caspase-3/7 activity in cardiomyocytes. The cells were extracted by enzyme digestion and inoculated into 96-well plates. The cells were transfected with 100nM miR-874 antagomiR or negative control. After 24 hours, the number of cardiomyocytes was detected by CellTiter-Fluor™ Cell Viability Assay as the internal reference. The Caspase-Glo ® 3/7 Assay was used to detect cardiomyocyte apoptosis.

#### Effects of miR-874 antagomiR on Proliferation of Cardiac Fibroblasts

Cardiac fibroblasts proliferation can secrete excess extracellular matrix leading to myocardial fibrosis, so we used CellTiter-Fluor™ Cell Viability Assay to detect the proliferation of cardiac fibroblasts. The results showed that the number of cardiac fibroblasts in the miR-874 antagomiR group did not change significantly (P=0.45> 0.05) compared with the negative control group (Fig.4). As a result, it can be seen that miR-874 antagomiR has no significant effect on the proliferation of cardiac fibroblasts.

**Fig. 4.**
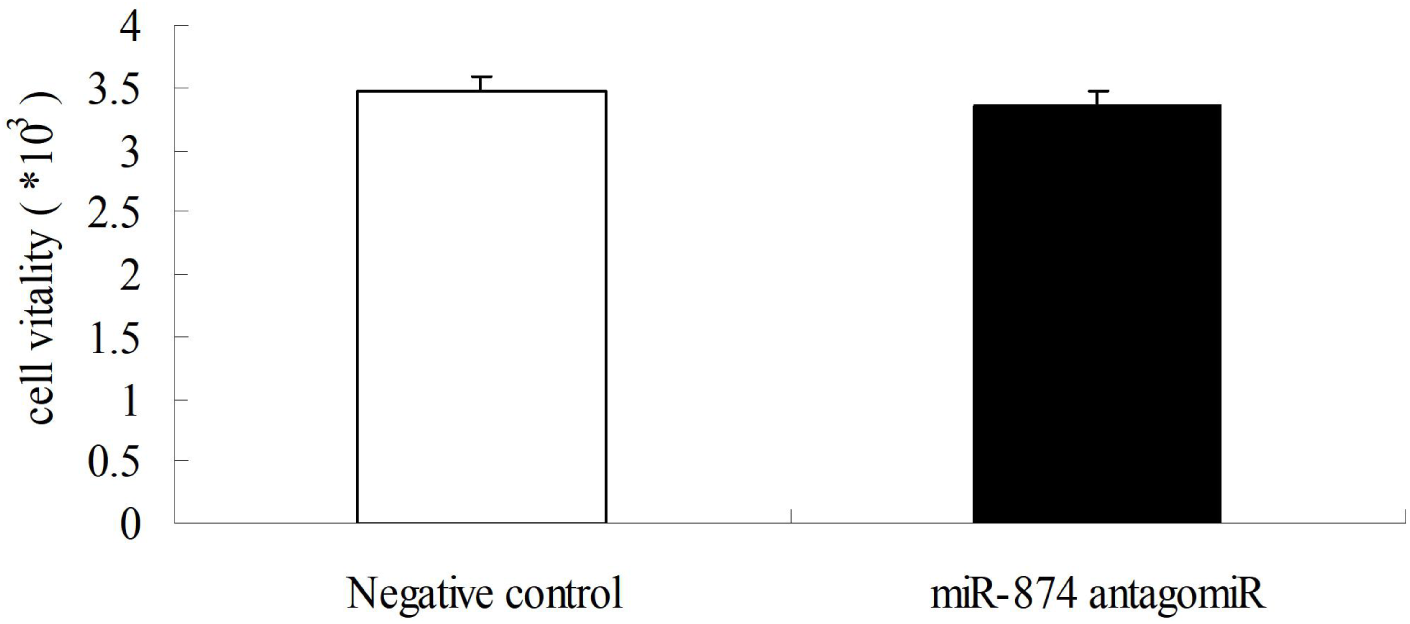
The proliferation of cardiac fibroblasts transfected with miR-874 antagomiR was detected using CellTiter-Fluor™ Cell Viability Assay. After cardiac fibroblasts were isolated by differential adhesion, the cells were plated into 96-well plates. Then the cells were transfected with 100nM miR-874 antagomiR or negative control, and the cell viability of cardiac fibroblasts was measured with CellTiter-Fluor™Assay.

### 3.4. miR-874 antagomiR Inhibits Cardiac Fibroblasts Apoptosis

The number of cardiac fibroblasts was examined by the CellTiter-Fluor™ Cell Viability Assay Kit as an internal reference, and the Caspase-Glo® 8 Assay was used to detect the apoptosis of cardiac fibroblasts. Fig.5 shows that the normalized caspase-8 activity of the miR-874 antagomiR group is significantly lower than negative control group (P=0.001<0.01). It was found that miR-874 antagomiR could inhibit cardiac fibroblasts apoptosis.

**Fig. 5.**
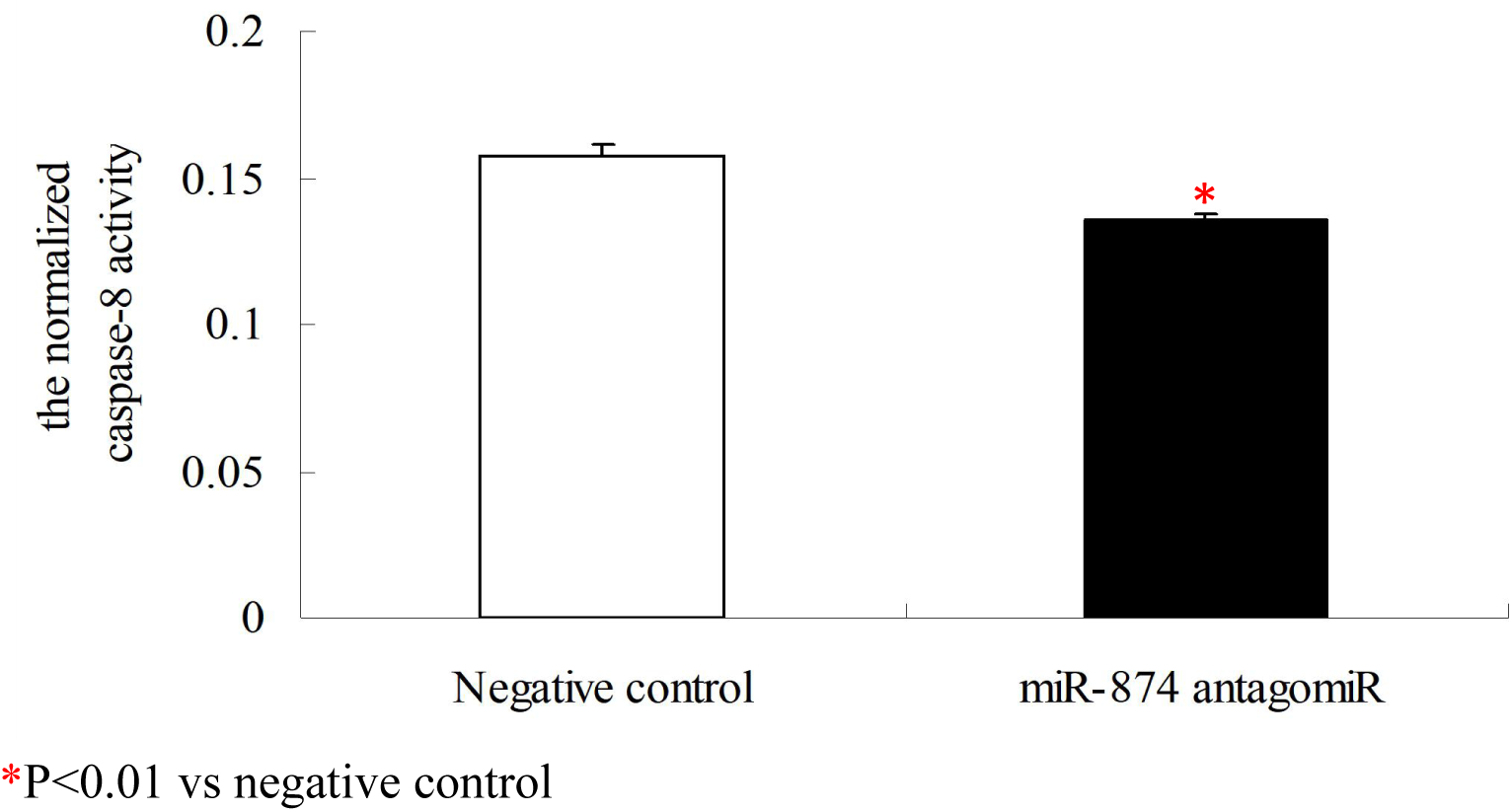
miR-874 antagomiR inhibits the normalized caspase-8 activities in cardiac fibroblasts. After cardiac fibroblasts were isolated by differential adhesion, the cells were seeded into 96-well plates. The 100nM miR-874 antagomiR or negative control was transfected into cells. The number of cardiac fibroblasts was measured with CellTiter-Fluor™Cell Viability Assay as the internal reference, and then Caspase-Glo® 3/7 Assay was used to detect the apoptosis of cardiac fibroblasts.*P<0.01 vs negative control.

### 3.5. miR-874 antagomiR Regulates the mRNA Levels of MMP2

The MMP2 and MMP9 can degrade extracellular matrix and affect the progress of myocardial fibrosis. Therefore, we examined the effect of miR-874 antagomiR on MMP2 and MMP9 mRNA levels in H_9_C_2_ cells. The expression levels of MMP2 and MMP9 relative to GAPDH were measured by qRT-PCR. The results of Fig.6 show that there was no statistically significant difference in MMP2 mRNA levels between the miR-874 antagomiR group and the negative control group (P=0.55>0.05), while the level of MMP9 mRNA in the miR-874 antagomiR group was lower than the negative control group (P=0.01<0.05). The results showed that miR-874 antagomiR could down-regulate the MMP9 mRNA level in H9C2 cells and had no effect on MMP2 mRNA level.

**Fig. 6.**
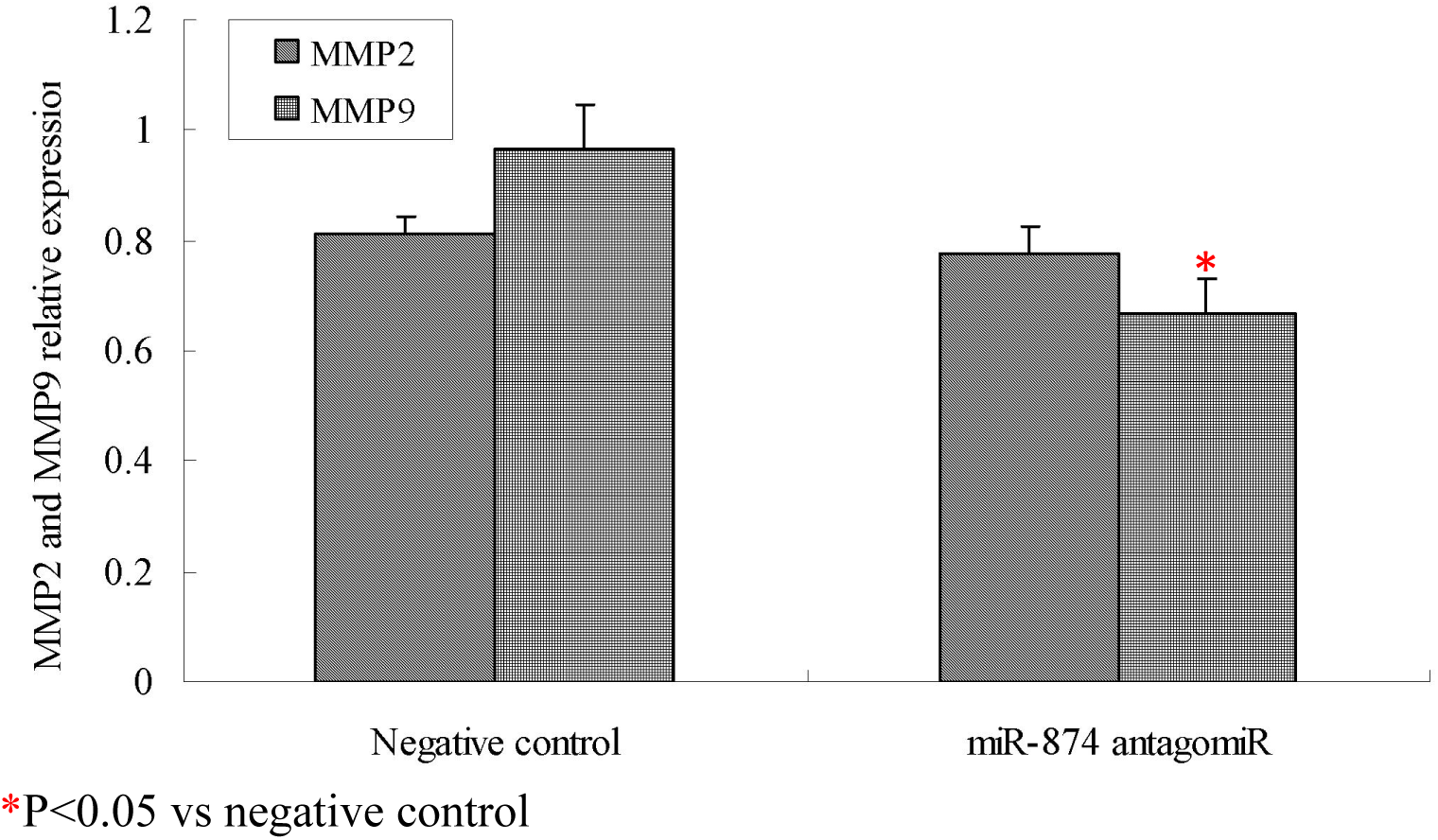
MMP2 and MMP9 expression was quantified by quantitative real time PCR. The H_9_C_2_ cells were plated into 60mm culture dish, transfected with 100nM miR-874 antagomiR or negative control and the collection and lysis of cells after 48h. Trizol was used to extract total RNA and reversely transcribed into cDNA, and then the mRNA levels of MMP2 and MMP9 were measured by qRT-PCR.*P<0.05 vs negative control

### 3.6. Effects of miR-874 antagomiR on SERCA2a mRNA and Ca^2+^ Concentration

The SERCA2a is responsible for the uptake of Ca^2+^ in the sarcoplasmic reticulum of the myocardium. The level of SERCA2a significantly affects myocardial sarcoplasmic reticulum Ca^2+^ uptake and overall myocardial contractility. The expression levels of SERCA2a relative to Dmd were detected by qRT-PCR. The results in Fig.7 show no significant difference in the level of SERCA2a mRNA between the miR-874 antagomiR group and the negative control group (P=0.48>0.05). This indicates that miR-874 antagomiR had no effect on SERCA2a mRNA levels in H_9_C_2_ cells.

**Fig. 7.**
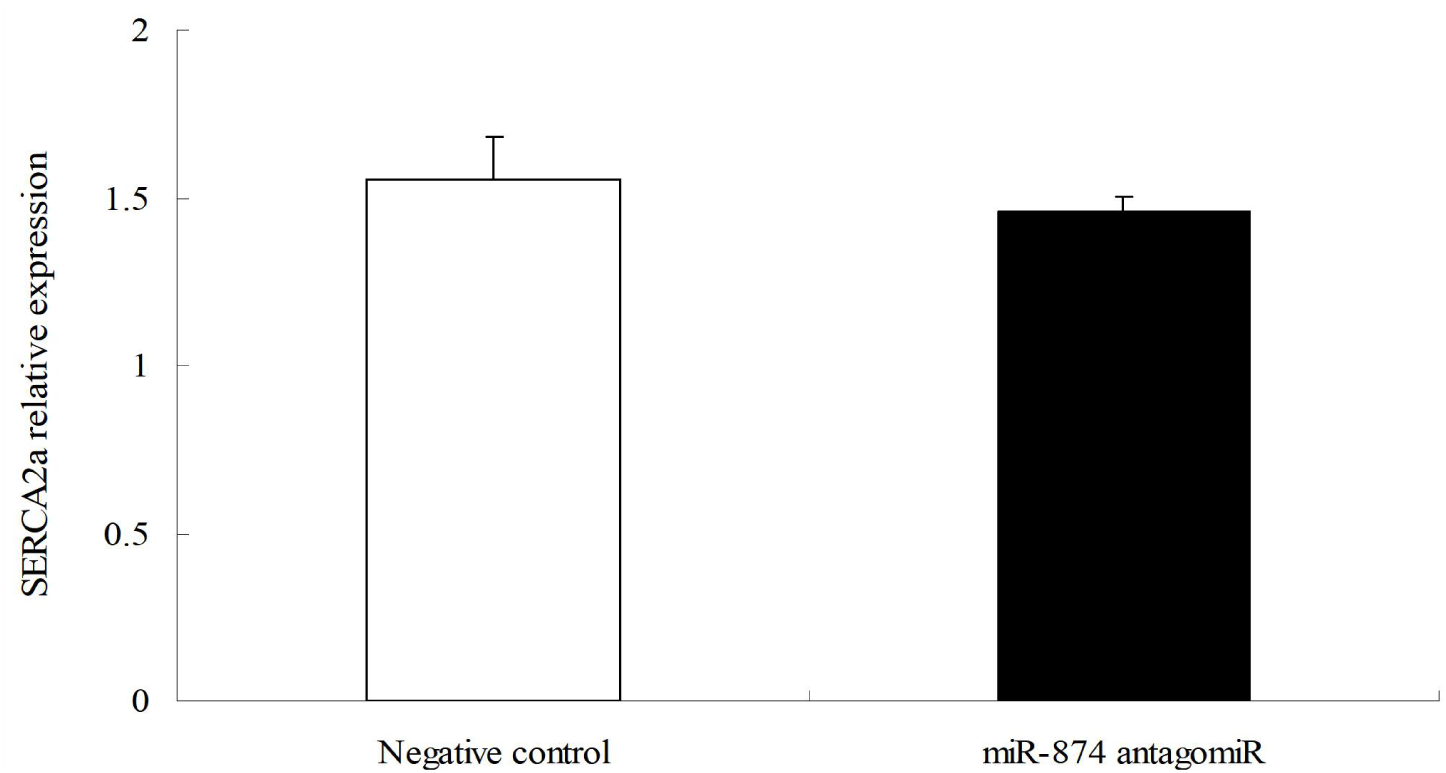
SERCA2a expression was quantified by quantitative real time PCR. H_9_C_2_ cells were plated into 60mm culture dishes and transfected with 100nM miR-874 antagomiR or negative control. After 48h, the total RNA was extracted with Trizol and reversely transcribed into cDNA. The mRNA level of SERCA2a was detected by qRT-PCR in cells.

Cardiac myocytes isolated from neonatal rats were seeded in 96-well plate and the beating cardiomyocytes in good condition were transfected with 100nM miR-874 antagomiR or negative control miRNA, miR-146a mimics, or mimics control respectively. The intracellular Ca^2+^ concentrations were measured by Fluo-4 NW Calcium Assay Kits according to the supplier’s instructions (Invitrogen) 48h after the transfection. Briefly, the growth medium from the adherent cell cultures was removed in order to eliminate sources of baseline fluorescence, particularly esterase activity. Then, they were washed with Hanks’ balanced salt solution and incubated with the Fluo-4 dye loading solution in the dark for 30 min at 37 C, The cells were then visualized and the calcium ion concentration was quantified with cytation 5 (Bioteck) which combines automated wide field microscopy with conventional multi-mode reading. Ca^2+^ fluorescence [excitation/emission wavelengths (EX/EM): 494/516 nm] was induced and recorded operating at ∼ 50 fps (frame per second). Then Ca^2+^ concentrations ([Ca^2+^]i) was calculated as follows: [Ca^2+^]i=Kd [F–Fmin]/[Fmax–F], where the Kd for Fluo-4/AM is 360 nmol/L, F is the mean fluorescent intensity of the cells, Fmin is the fluorescent intensity when the cells were treated with 5 mmol/L EGTA (Sigma, USA), and Fmax is the fluorescent intensity when the cells were treated with 0.1% Triton X-100 (Sigma, USA).

Cardiomyocytes were incubated with Ca^2+^ fluorescent indicator Fluo-4 NM and 10% Triton x-100 and 10nM EGTA, and fluorescence (F, Fmax and Fmin) was detected at 494_EX_/516_EM_. F, Fmax and Fmin were used to calculate Ca^2+^ concentration in cardiomyocytes. Two groups of cardiomyocytes were incubated with the Ca^2+^ fluorescent indicator Fluo-4 NM, which showed that fluorescence was produced by the combination of Fluo-4 NM with Ca^2+^ in the cells (Fig.8a-8b). As shown in Fig.8, in the positive control group, the concentration of Ca^2+^ in the cardiomyocytes transfected with miR-146a mimics was significantly lower than the mimic negative control group (P=0.03<0.05). There was no statistically significant difference in intracellular Ca^2+^ concentration between the miR-874 antagomiR group and the antagomiR negative control group (P=0.88>0.05) as shown in Fig.9.

**Fig. 8.**
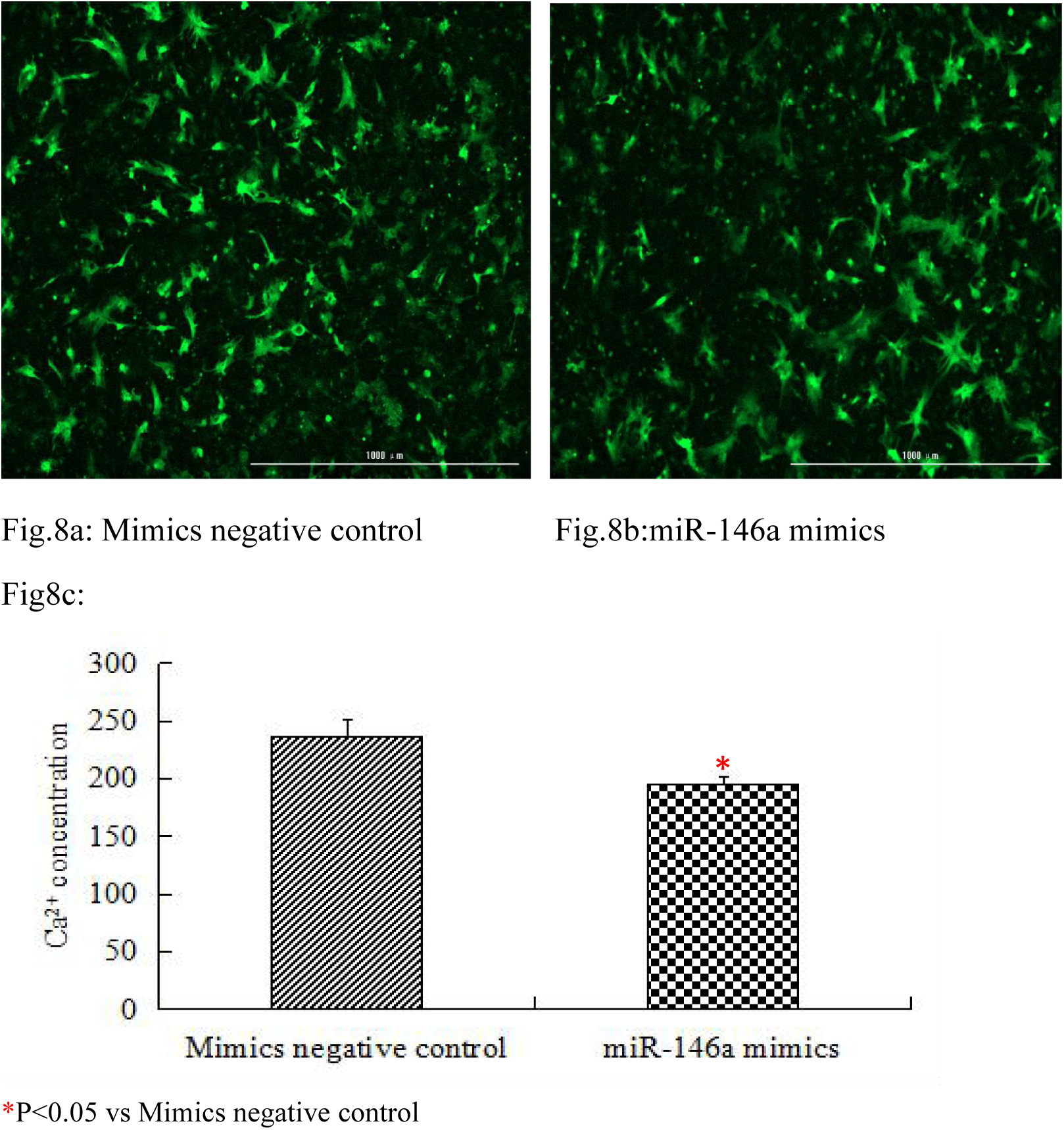
The miR-146a mimics inhibits calcium concentration in Cardiomyocytes. Cardiomyocytes were extracted by enzyme digestion and inoculated into 96-well plates.The cells were transfected with miR-146a mimics or mimics control. After 72 h, the concentration of Ca^2+^ in cardiomyocytes was measured using Fluo-4 NW Calcium Assay Kits.

**Fig. 9.**
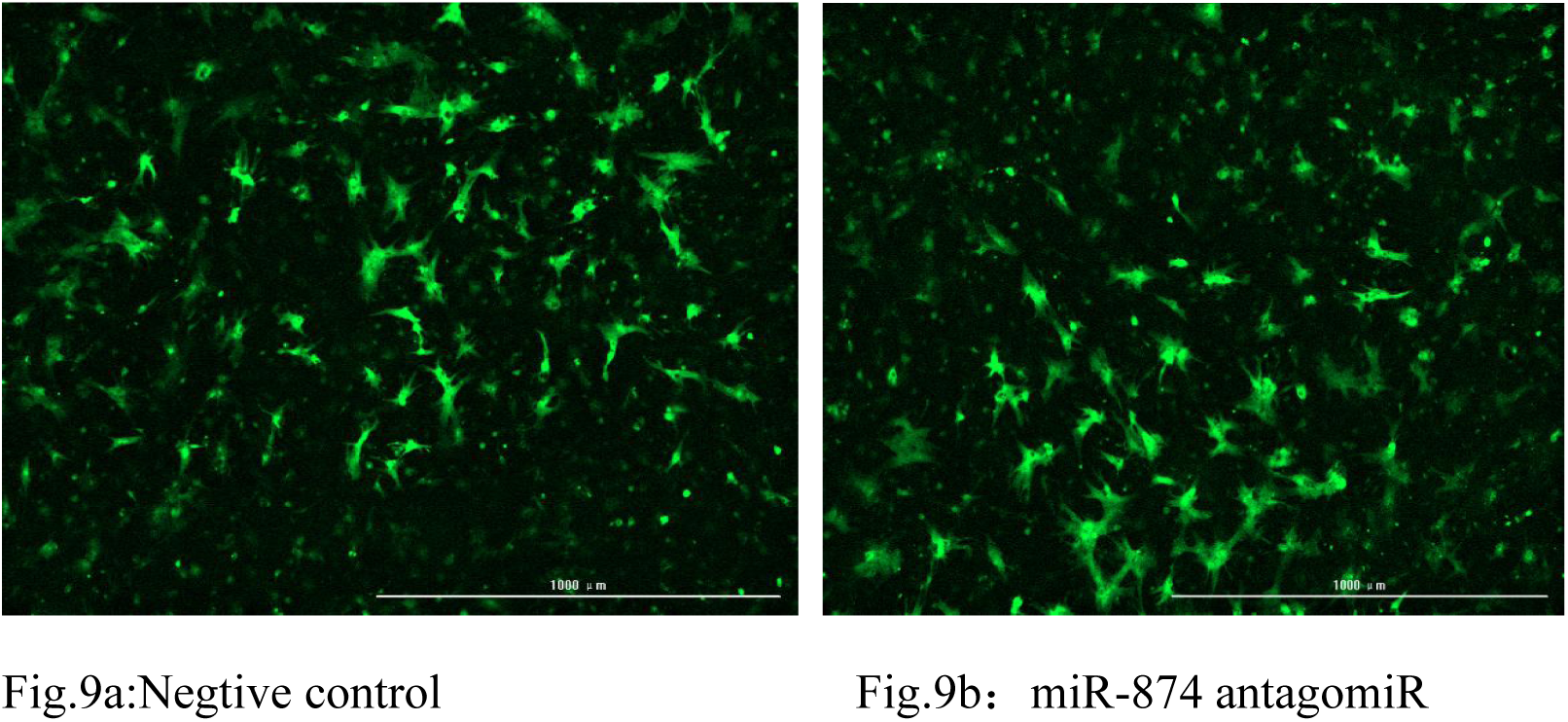
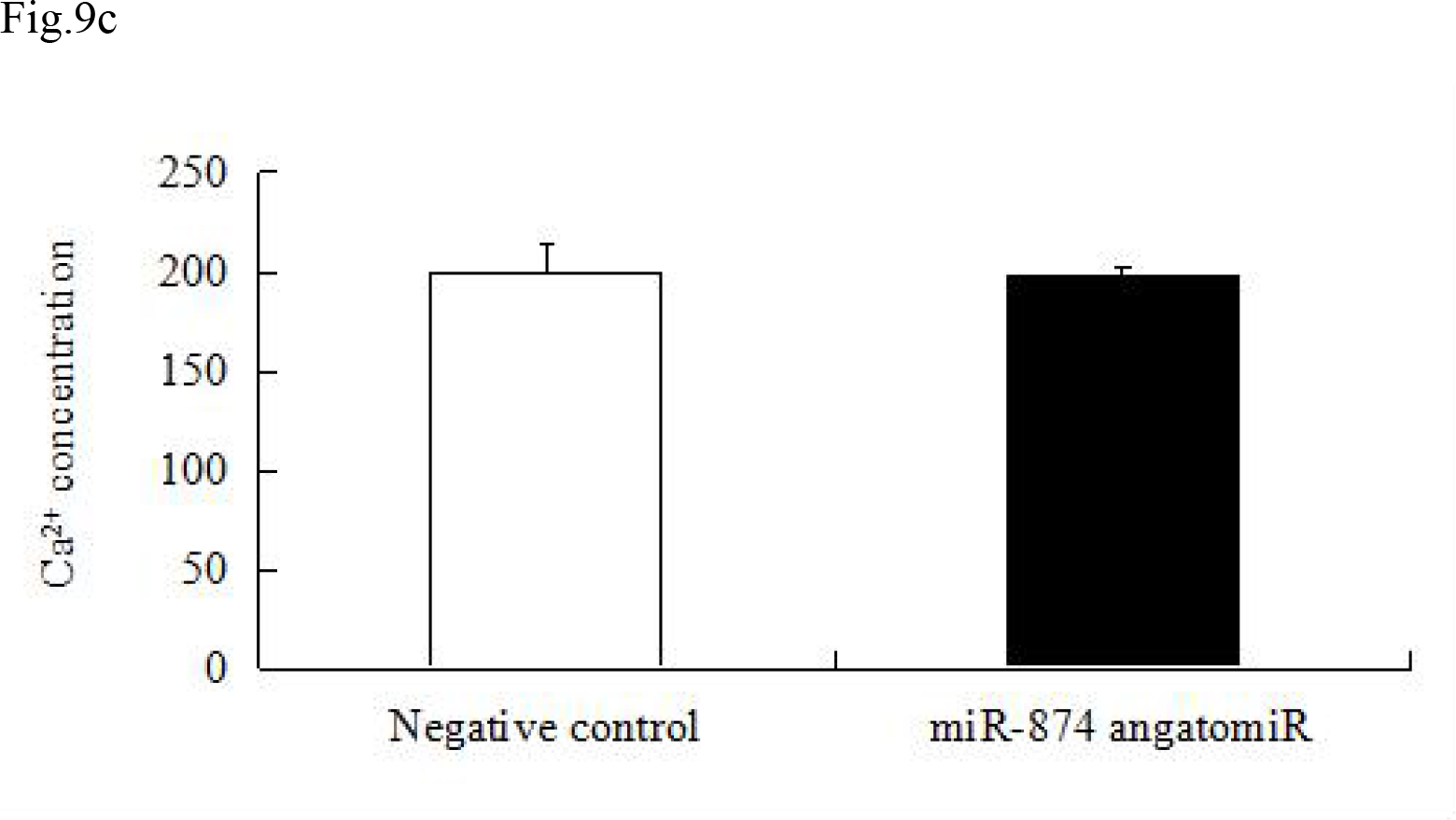
The effect of miR-874 angatomiR on calcium concentration in Cardiomyocytes. Cardiomyocytes were extracted by enzyme digestion and seeded into 96-well plates. The cells were transfected with either miR-874 angatomiR or negative control, respectively. After 72 h, the concentration of Ca^2+^ in cardiomyocytes was measured using Fluo-4 NW Calcium Assay Kits.

## 4. Discussion

Cardiovascular diseases are the leading cause of death globally. Heart failure is the end stage of cardiovascular diseases caused by multiple factors. Death of cardiomyocytes is an important cause for the development of heart failure and myocardial infraction. MicroRNAs can participate in many important biological processes such as cell growth, differentiation, necrosis, as well as proliferation. Many studies have found that miRNAs exerted significant regulatory functions in heart failure,myocardial infraction, arrhythmias, hypertension, and many other diseases. To date, most studies of miR-874 focused on cancer, and their findings suggest that miR-874 could inhibit cancer cell proliferation, metastasis and invasion, but regarding to the treatment of heart failure, further studies are needed to understand the relationship between miR-874 and cardiomyocytes. Using the biochip technology, a recent study detected significantly increased expression of miR-874 in cells treated with hydrogen peroxide. Thus, to use miR-874 as an intervention strategy might provide new insights for treating myocardial necrosis. Studying the relationship between miR-874 inhibitors and myocardial necrosis is of significance for the development of heart failure drugs, thus this study aimed to discuss the effects of miR-874 inhibition on caspase-8 expression and rat H_9_C_2_ cell proliferation.

Cardiomyocyte death is an important cause of heart failure (Konstantinidis K et al., 2012), and there are two forms of cardiomyocyte death: necrosis and apoptosis. Cell apoptosis is described as a gene-controlled, autonomous, programmed cell death to maintain the stability of the internal environment, it is not a phenomenon of self-injury under pathological conditions, but to better adapt to the living environment and actively fight for a death process, it is also known as programmed cell death. While necrosis is a passive death caused by pathological conditions, it is characterized by swelling of cells, dissolution of organelles and rupture of cell membranes, accompanied by tissue inflammation (Kroemer G et al., 2005). Studies have found that cardiomyocyte apoptosis and heart failure have a certain connection (Gaballa MA et al., 2002). Cardiomyocytes are terminally differentiated cells that no longer undergo cell division shortly after birth. Loss of cardiomyocytes can cause pathological damage to the heart, and may cause heart failure. Caspase belongs to a cysteine protease family, usually present in the cells in the form of zymogen (Stepien A et al., 2007). It is widely accepted that caspase-3 is the key protease inducing apoptosis, once activated, the apoptosis will occur inevitably, hence the caspase-3 is all called “death caspase”. Caspase is divided into start-up type and effect type, in which caspase-3 and caspase-8 are at the core position in initiating and performing apoptosis (Moe GW et al., 2016). At the same time, studies have shown that in Hela cells by quercetin-induced apoptosis, caspase-3 and caspase-8 activities are significantly higher than the control group (Granado-Serrano AB et al., 2006; Chao G et al., 2011). We detected caspase-3 activities in cardiomyocytes by Caspase-Glo® 3/7Assay, Results showed that miR-874 antagomiR has no effect the activities of caspase-3.

There are many causes for heart failure and myocardial fibrosis is one of them. Myocardial fibrosis is caused by extracellular matrix deposition while the synthesis of myocardial collagen is greater than degradation. MMPs play a key role in the maintenance of normal myocardial extracellular matrix decomposition and synthesis in myocardial reconstruction (Sivasubramanian N et al., 2001; Yamazaki T et al., 2004; Cogni AL et al., 2013). Cardiac fibroblasts secrete excess extracellular matrix to form scar tissue, and lead to high expression of contractile protein alpha smooth muscle actin (αSMA), altering the surrounding extracellular matrix (Leask A., 2015). It is well known that angiotensin II can induce cardiac fibrosis.

Ma Yue et al used AngII to induce cardiomyocytes fibrosis and then cardiac fibroblasts were labelled with Edu and displayed red fluorescence. It was found that the number of cardiac fibroblasts increased significantly in AngII treated cells (Li YQ et al., 2013; Yue M et al., 2013).

We used CellTiter-Fluor ™Cell Viability Assay and Caspase-Glo® 8 Assay to detected cardiac fibroblast proliferation and apoptosis, and found there was no significant difference in proliferation between the miR-874 antagomiR transfected cells and scramble control transfected cells; however, there was significant difference between the caspase 8 activity between the the miR-874 antagomiR transfected cells and scramble control transfected cells. It suggests that miR-874 antagomiR might suppress the apoptosis of cardiac fibroblasts.

Cardiac fibrosis involves the excessive accumulation of extracellular matrix (ECM) in the heart, which leads to cardiac dysfunction, and is closely associated with numerous cardiovascular diseases, including hypertension, myocardial infarction and cardiomyopathy. As the most common cell type in the heart, cardiac fibroblasts (CFs) play a pivotal role in the development of cardiac fibrosis via the excessive synthesis of collagens and the degradation of ECM via the production of matrix metalloproteinases (MMPs). The MMP family is considered to be profoundly involved in the pathogenesis of cardiac fibrosis and plays a crucial role in cardiac remodeling.

Ang II increases collagen expression, proliferation, and migration in CFs by activating a variety of cell signaling pathways such as transforming growth factor β (TGF-β) and mitogen-activated protein kinases (MAPKs) pathways, which promote the differentiation, proliferation and migration of CFs. Cardiac fibrosis can be induced by angiotensin II, it is successful to use AngII to establish model of cell proliferation and collagen synthesis (Ruoyu L et al., 2010). Therefore, the proliferation and apoptosis of cardiac fibroblasts affect the degree of myocardial fibrosis. Our study examined the proliferation and apoptosis of cardiac fibroblasts. The results showed that there was no significant difference in the proliferation of cardiac fibroblasts between the miR-874 antagomiR transfected cells and the negative control transfected cells; however the apoptosis induced by miR-874 antagomiR in two groups was significantly different. The miR-874 antagomiR significantly inhibited the apoptosis of cardiac fibroblasts, thus increasing myocardial fibrosis and affect heart failure.

Matrix metalloproteinases (MMPs) belong to the zinc ion (Zn^2+^) -dependent endopeptidase family, which specifically degrades myocardial extracellular matrix (ECM) and thus affecting the occurrence and progression of heart failure (Lee RT et al., 2002; Rai V et al., 2016). In the development of heart failure, the activity and content of MMP2 and MMP9 show an increasing trend (Han CK et al., 2017; Chegeni S et al., 2015). In the patients with cardiac hypertrophy, collagen volume fraction (ICVF) and perivascular collagen area (PVCA), and the mRNA levels of type?and type ? collagen in myocardium are higher than those in control group, while the expression of MMP2 and MMP9 are significantly increased (Xianmei W et al., 2006).

In the animal model of myocardial infarction, knockout of the MMP9 gene causes a decrease in myocardial infarct size and collagen deposition in the infarct area and left ventricular dilatation, indicating that MMP9 plays an important role in ventricular remodeling after infarction (Ducharme A et al., 2000; Romanic AM et al., 2002). Our study used qRT-PCR to detect the mRNA levels of MMP2 and MMP9 in H_9_C_2_ cells. The results showed that miR-874 antagomiR had no effect on MMP2 mRNA level but it could inhibit the mRNA level of MMP9. Therefore, miR-874 antagomiR can relieve myocardial fibrosis by inhibiting the mRNA level of MMP9 in cells.

Cardiac fibrosis and cardiomyocyte apoptosis leads to the progression to heart failure. Cardiomyocyte cell death is an important factor in the pathogenesis of conditions such as heart failure and cardiac infarction. Cardiomyocyte apoptosis is one form of cell death during heart failure. Moreover, the contraction of cardiomyocyte is triggered by calcium release. In cardiac muscle, the contraction and relaxation rhythm is tightly associated with the uptake and release of calcium by the sarcoplasmic reticulum. Calcium overload leads to cell injury and play an important role in the deterioration of cardiac function.

Ca^2+^ participates in the cardiac function and is an important factor of heart muscle excitation-contraction (E-C) coupling (Arai M et al., 1994). Calcium (Ca^2+^) plays an important role in the signal transduction pathways as a ubiquitous messenger in cardiac myocytes, thus disruption of the intracellular Ca^2+^ homeostasis and transportation results in the cardiac function disabilities, heart failure, cardiac injury, and infarction SERCA2a is called sarcoplasmic reticulum Ca^2+^-ATPase, and SERCA2a is a subtype mainly present in cardiomyocytes. In recent years, more and more studies are about treatment of heart failure through vector mediated SERCA2a overexpression. In the animal model of heart failure, SERCA2a overexpression in animal myocardium is mediated by recombinant adeno-associated virus (rAAV). The results show that SERCA2a expression is significantly higher than the heart failure group, and effectively improved hemodynamics in the animal of heart failure, enhanced systolic and diastolic function of heart, increased cardiac energy metabolism, the severity of heart failure declined, reduced myocardial necrosis and delayed ventricular remodeling, it can prevent arrhythmia after myocardial infarction (Li XY et al., 2006; Mi YF et al., 2009; Wei X et al., 2011). In the model of chronic heart failure established by combined aortic valve disruption and abdominal aortic stenosis, the activity and mRNA level and protein expression of SERCA2a are lower than sham operation group, and at the same time, the calcium intake capacity of sarcoplasmic reticulum decreased (Periasamy et al., 2001)2. A number of studies have shown that the Ca^2+^ in cardiomyocytes is loaded with Fluo-3 AM fluorescent indicator, and the results show a significant increase in Ca^2+^ concentration in animal heart failure models (Li Y et al., 2009; Zhang X et al., 2009; Menglu M et al., 2013). The Ca^2+^ is involved in cardiac function, which is not only an important factor in the excitation contraction coupling of the myocardium, but also plays an important role in the signal transduction of cardiomyocytes; The calcium steady state abnormalities and transport disorders in the myocardium, it will cause myocardial dysfunction, heart failure and myocardial injury, infarction, etc. (Chakraborti S et al., 2007; Kubalova Z et al., 2005; Zima AV et al., 2006). Therefore, the SERCA2a abnormalities in cardiomyocytes, leading to calcium overload will damage the cardiomyocytes, and then develop to heart failure.

In myocardium, SERCA2a is the major protein participating in the regulation of calcium and it plays a key role in the myocyte calcium cycling and contraction and relaxation cycling.

Human and animal heart failure experiments in recent years have demonstrated that expression and activity of the SERCA2a protein are significantly reduced in the heart failure myocytes than the normal myocytes. And we found miR-146a regulated the calcium concentration in neonatal cardiomyocytes.

In our study, qRT-PCR was used to detect SERCA2a mRNA levels in H_9_C_2_ cells, and Ca^2+^ concentrations were measured using Fluo-4 NM in cardiomyocytes. The results showed that SERCA2a mRNA and Ca^2+^ concentration are not different between the miR-874 antagomiR group and the negative control group in cells. From this point of view, miR-874 antagomiR may have no effect on the treatment of heart failure.

In summary, miR-874 antagomiR and miR-146a have both advantages and disadvantages in the treatment of heart failure. Therefore, we need to further investigate the relationship between miR-874 antagomiR or miR-146a and heart failure, which will provide a basis for future development of heart failure treatment drugs.

## Acknowledgments

This project was supported by the grants from the National Natural Science Foundation of China (No. 81070128); Postgraduate Scientific and Technological Innovation Foundation of Shenyang Medical College (Y20170609); Shenyang “High-level Innovative Talent Program”(RC170408); Liaoning Provincial Key R&D Program (2018225013); Shen Yang Young high-level talents Program (RC180379); National Innovation & Entrepreneurship Training Program for College Students (201810164005); Distinguished Young Scholar Cultivation Plan of Liaoning (Excellent Talent Program by Provincial Education Department);

